# Reproduction in deep-sea vent shrimps is shaped by diet and phylogeny, with rhythms unlinked to surface production

**DOI:** 10.1101/2021.09.10.456763

**Authors:** Pierre Methou, Chong Chen, Hiromi K. Watanabe, Marie-Anne Cambon-Bonavita, Florence Pradillon

**Affiliations:** Univ Brest, Ifremer, CNRS, Laboratoire de Microbiologie des Environnements Extrêmes, UMR6197, 29280 Plouzané, France; Ifremer, Centre Brest, Laboratoire Environnements Profonds (REM/EEP/LEP), 1625 route de Sainte Anne, 29280 Plouzané, France; X-STAR, Japan Agency for Marine-Earth Science and Technology (JAMSTEC), Yokosuka, 237-0061, Japan

**Keywords:** Biological rhythms, crustacean, deep sea, hydrothermal vent, reproduction, seasonality, trophic ecology

## Abstract

Variations in offspring production according to feeding strategies or food supply have been recognized in many animals from various ecosystems. Despite an unusual trophic structure based on non-photosynthetic primary production, these relationships remain largely under-studied in chemosynthetic ecosystems. Here, we use *Rimicaris* shrimps from deep-sea hydrothermal vents as a study case to explore relations between reproduction, diets and food supply in these environments. For that, we compared reproductive outputs of three congeneric shrimps differing by their diets. They inhabit vents located under oligotrophic waters of tropical gyres with opposed latitudes, allowing us to also examine the prevalence of phylogenetic vs environmental drivers in their reproductive rhythms. For this we used both our original data and a compilation of published observations on the presence of ovigerous females covering various seasons over the past 35 years. We report distinct egg production trends between *Rimicaris* species relying solely on chemosymbiosis – *R. exoculata* and *R. kairei* – and those relying on mixotrophy – *R. chacei* – where *R. chacei* produces broods with higher numbers of smaller eggs. Besides, our data and historical records suggest a reproductive period with substantial proportions of brooding females mostly between January and early April for all examined species whatever the region. Intriguingly, this periodicity does not correspond to seasonal variations in surface production with presence of brooding females during either boreal winter or austral summer. These observations contrast with the long-standing paradigm in deep-sea species for which periodic reproductive patterns have always been attributed to seasonal variations of photosynthetic production sinking from surface. Our results suggest the presence of intrinsic basis for biological rhythms in the deep sea, and bring to light the importance of having year-round observations in order to understand life history of vent animals.

## Introduction

Relationships between feeding strategies and offspring’s production have been extensively studied for a number of animal phyla in various ecosystems (Pierotti and Annett 1990, Qian and Chia 1991, Broderick et al. 2003, Meiri et al. 2012, Sibly et al. 2012). Where no clear trends between trophic regime and reproductive outputs could be observed in lizards (Meiri et al. 2012), large variations of fecundity or reproductive outputs according to diets have been highlighted in others including birds (Pierotti and Annett 1990, Sibly et al. 2012), sea turtles (Broderick et al. 2003), annelids (Qian and Chia 1991), insects (Gutiérrez et al. 2020), or crustaceans (Lin and Shi 2002, Tziouveli et al. 2011, Almeida et al. 2018). All these studies are based on ecosystems where photosynthetic primary production is the sole basis of food webs. A number of ecosystems on Earth, such as deep-sea hydrothermal vents, cold seeps and organic falls, rely on another primary organic source: the chemosynthesis (German et al. 2011). In chemosynthetic-based ecosystems lush and dense faunal communities are supported by microbial communities using chemical energy arising from geofluids to produce their organic matter, and animals there often form intricate symbiotic associations with these chemosynthetic bacteria for nutrition (Dubilier et al. 2008). From total reliance on symbiosis to bacterivory to mixotrophy to carnivorous, these particular ecosystems display a variety of diets in networks often differing strikingly from those found elsewhere (Govenar 2012). In this context, these environments are natural laboratories to study relationships between reproduction and trophic regimes. However, no study has yet explored these relationships extensively in chemosynthetic-based ecosystems, with variations in the reproductive output among or within animal groups being mostly attributed to phylogenetic constraints (Tyler and Young 1999) or variations in female body size (Ramirez-Llodra et al. 2000, Copley and Young 2006, Nye et al. 2013, Marsh et al. 2015, Marticorena et al. 2020).

Although most species from deep-sea chemosynthetic-based ecosystems exhibit continuous reproductive patterns (Jollivet et al. 2000, Faure et al. 2007, Tyler et al. 2008, Hilário et al. 2009, Marticorena et al. 2020), seasonal cycles following variations in surface primary production have been observed in many taxa such as bythograeid crabs, alvinocaridid shrimps, and bathymodiolin mussels (Perovich et al. 2003, Dixon et al. 2006, Copley and Young 2006, Tyler et al. 2007). In some groups, the surface production pattern is clearly linked to reproductive patterns in the deep. For example, some vent crab species exhibit continuous gametogenesis under the highly oligotrophic waters of the South Pacific Gyre, while others living in more productive areas north of this gyre have seasonal reproduction (Perovich et al. 2003, Hilário et al. 2009). Such examples have led to the paradigm that rhythms of sinking photosynthetic matter is the key time-synchronising cue for deep-water animals with seasonal reproduction, signalling individuals to spawn at the same period of the year.

The symbiotic and sister-species shrimps *Rimicaris exoculata* and *R. kairei* are among the most iconic hot-vent animals, with unique adaptations such as hosting symbiotic bacteria in their enlarged head cavity and the fused “dorsal eye”, a novel sensory system representing a pinnacle of adaptation to vents (Zbinden and Cambon-Bonavita 2020). The entire family they belong to, Alvinocarididae, is endemic to vents and seeps with different taxa exhibiting various trophic regimes and various levels of reliance on symbiosis (Rieley et al. 1999, Gebruk et al. 2000, Apremont et al. 2018), constituting an ideal case for elucidating relationships between reproduction and diet in chemosynthesis-based ecosystems. Species with different trophic regimes often co-occur on same vent fields, for example *R. exoculata* which rely on symbiosis and the mixotrophic *R. chacei,* allowing for a straight-forward comparison of species. The symbiotrophic species such as *R. exoculata* and *R. kairei* exhibit greatly inflated cephalothorax and live in large aggregations around the hot fluid flows (Rieley et al. 1999, Gebruk et al. 2000, Van Dover 2002, Streit et al. 2015, Methou et al. 2020), whereas mixotrophic species such as *R. chacei* (Gebruk et al. 2000, Apremont et al. 2018, Methou et al. 2020) lacks the inflated cephalothorax and lives in small groups (Methou et al. 2021).

Attempts to understand the reproduction of *Rimicaris* shrimps have so far faced contradicting evidences. While continuous reproduction was initially suggested from examination of their reproductive tissues (Ramirez-Llodra et al. 2000, Copley et al. 2007) hardly any ovigerous individuals could be collected in over 35 years of seagoing expeditions, despite a relatively focused and repeated sampling effort (Williams and Rona 1986, Vereshchaka 1997, Shank et al. 1998, Ramirez-Llodra et al. 2000, Copley et al. 2007, Komai and Segonzac 2008). Brooding females were only found in large numbers in 2014 (Methou et al. 2019, Hernández-Ávila et al. 2021) and to a lesser extent in 2007 (Guri et al. 2012), around a restricted period between January and March. These females were living within dense aggregations close to vent orifices (Hernández-Ávila et al. 2021), refuting previous hypothesis that brooding females would migrate to vent periphery to protect their eggs from vent fluid, as observed for bythograeid crabs (Perovich et al. 2003) as well as *Kiwa* squat lobsters (Marsh et al. 2015). Taken together, these recent findings suggest a seasonal reproductive cycle possibly linked to surface primary production, as shown in vent crabs, bathymodiolin mussels and some other alvinocaridid shrimps (Dixon et al. 2006, Copley and Young 2006, Tyler et al. 2007). However, the low primary productivity of oligotrophic surface waters and the overall low export to the deep-sea compartment within the region where these shrimps have been collected (−3500 m depth) (Pabortsava et al. 2017, Harms et al. 2021) challenges the possibility of such a link.

Here, we compare reproductive outputs – i.e., fecundities and egg volumes – of three congeneric vent shrimps with contrasting diets, using two species with a full nutritional dependence on their chemosymbiosis (i.e., *R. exoculata* and *R. kairei)* and one with a mixotrophic diet combining symbiosis, bacterivory, and scavenging (i.e., *R. chacei)* to find out if links between diets and reproduction are also present in chemosynthetic based ecosystems. Distribution of these species in two different hemispheres (Northern Atlantic and Southern Indian Ocean) with similar levels of, but seasonally opposed, surface production allows us to evaluate if their brooding periods are in synchrony with patterns of surface primary productivity.

## Materials & Methods

### Field sampling

Shrimps were collected from three hydrothermal vent fields (figure 1): TAG (26°08’2.12”N, 44°49’36.12”W, 3620 m depth) and Snake Pit (23°22’5.88”N 44°570”W, 3470 m depth) on the Mid Atlantic Ridge (MAR) between March and April in 2017 (HERMINE (doi:10.17600/17000200)) and between February and March in 2018 (BICOSE2 (doi:10.17600/18000004)) for *R. exoculata* and *R. chacei,* and Kairei (25°19’10.2”S, 70°2’24”E, 2415 m depth) on the Central Indian Ridge (CIR) between February and March both in 2002 and 2016, as well as in November 2009 for *R. kairei.* Shrimps were sampled using suction samplers of the human-occupied vehicle (HOV) *Nautile* or the HOV *Shinkai 6500.* Upon recovery on-board, ovigerous females of *R. exoculata* and *R. chacei* were identified and individually sorted from the rest of the population. An additional published dataset for *Rimicaris exoculata* collected at TAG and Snake Pit in January-February 2014 is also included in our analyses and available in the Ifremer SEANOE (SEA scieNtific Open data Edition) database at: https://doi.org/10.17882/84112.

**Figure 1.**
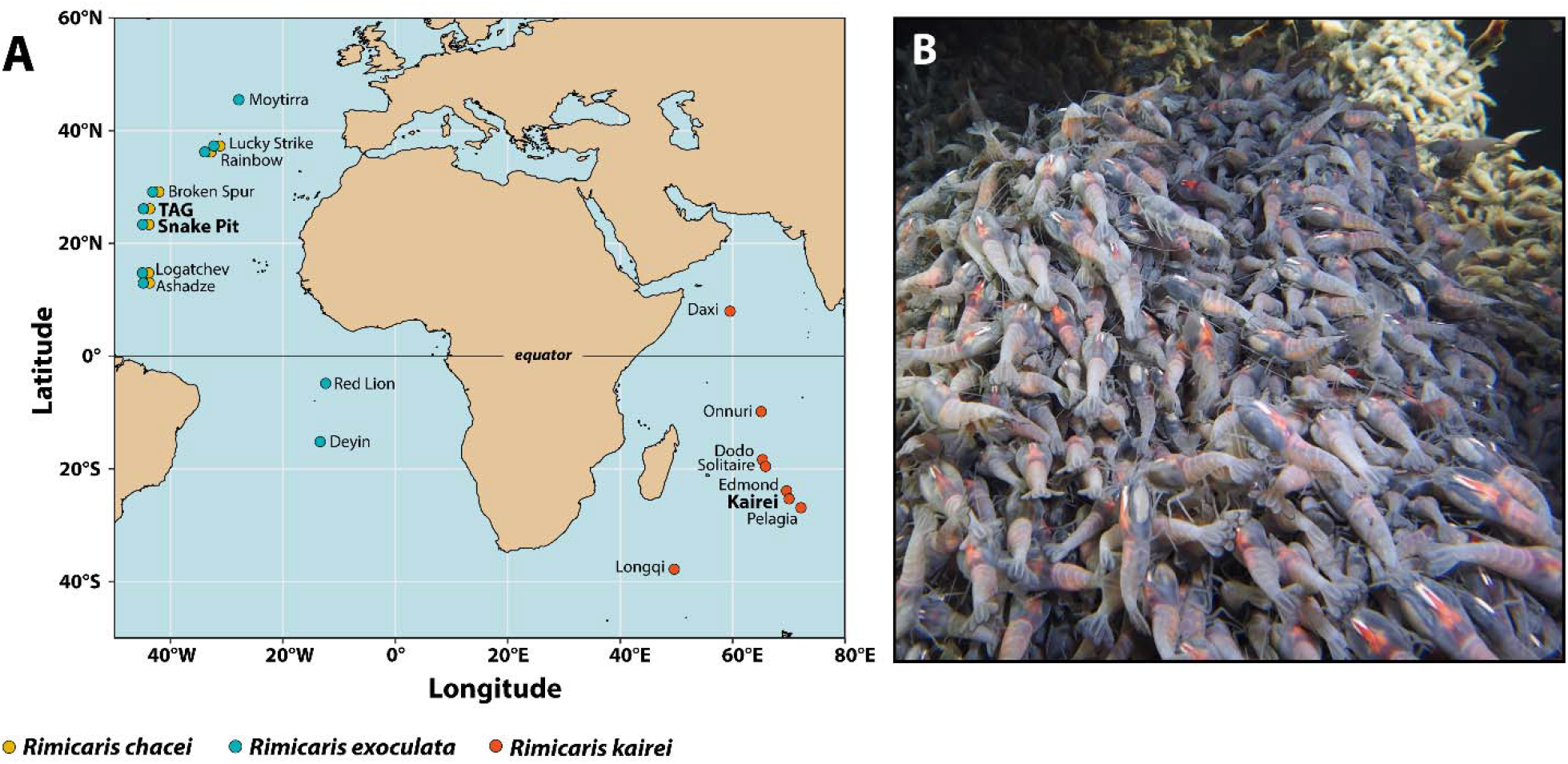
Context of the study, **a.** Current geographic distribution of the three *Rimicaris* species studied, **b**. Dense aggregations of *R. kairei* shrimps in February 2016 at the Kairei hydrothermal vent field.

Ovigerous females include both brooding females and females that have just released their larvae but still retained modified pleopods, characteristic of a recent brooding status, following Nye at al. 2013 (Nye et al. 2013). For *R. kairei,* ovigerous females were identified and sorted from the rest of the population in the laboratory after the cruise. Egg stages in broods were identified on-board during the BICOSE2 expedition before being fixed in 10% neutral buffered formalin for 24h and then rinsed and stored in 80% EtOH. For other expeditions, ovigerous females were fixed and stored in 10% formalin on-board and identification of egg stages was conducted in shore-based laboratory.

### Shrimps and Eggs Measurements

Carapace length (CL) of each female was measured with Vernier callipers from the posterior margin of the ocular shield (or eye socket) to mid-point of the posterior margin of the carapace, with an estimated precision of 0.1 mm. For each brood, total number of eggs was manually counted. Ten eggs were selected randomly to measure maximum and minimum diameters. Volume of embryos was estimated following the same method as (Hernández-Ávila et al. 2021), considering a spheroid volume v= (4/3).π.r1.r2^2^, where rl and r2 are half of maximum and minimum axis of each egg, respectively. Eggs in broods were classified in three developmental stages (early, mid, and late), following the classification used in (Methou et al. 2019). Some ovigerous females harboured damaged or aborted broods and were thus discarded from our fecundity analysis. Furthermore, we also compiled data for the presence of ovigerous female shrimps from the literature and using dive videos from cruises over the last 35 years, in order to assess the reproductive seasonality. Proportions of ovigerous females within samples were calculated against the number of sexually mature females (See supplementary material). Dataset compiling size, reproductive status, fecundities, egg developmental stages, and egg diameters of each shrimp individual is available in the Ifremer SEANOE (SEA scieNtific Open data Edition) database at: https://doi.org/10.17882/84195.

### Statistical analysis

Brooding females were grouped according to sampling year, species and vent field. Visual examination of our dataset and Shapiro-Wilk normality tests revealed that size, egg number, and egg volume of *Rimicaris* brooding females deviated from normal distribution. Therefore, non-parametric tests were used for intergroup comparison, with a Kruskal-Wallis test followed by post-hoc Dunn tests when three or more groups were compared. Spearman rank-order correlation were used to assess relationship between log_e_-transformed realized fecundity and log_*e*_-transformed carapace length of the different shrimp groups. Egg stage proportions between groups were compared using χ^2^ test with Yate’s correction. All analyses were performed in R v. 4.0.3 (R Core Team 2020).

## Results

### Interannual variation of *R. exoculata* reproduction

Realized fecundity of *Rimicaris exoculata* correlated positively with carapace length in both 2014 (Spearman correlation *r*= 0.67, *p* < 0.0001) and 2018 (Spearman correlation *r*= 0.73, *p*< 0.0001) (figure 2a). Moreover, slopes of the relationship between log_*e*_-transformed realized fecundity and log_*e*_-transformed carapace length did not differ significantly between these two sampling years (*F*= 1.175, *p* = 0.28) or between TAG and Snake Pit in 2018 (*F*= 0.004, *p*= 0.95). Although no correlation was found in 2017 (Spearman correlation *r*= 0.38, *p*= 0.16), probably due to the lower sampling effort, realized fecundity in 2017 still fell within the range of the other sampling years (figure 2a).

**Figure 2.**
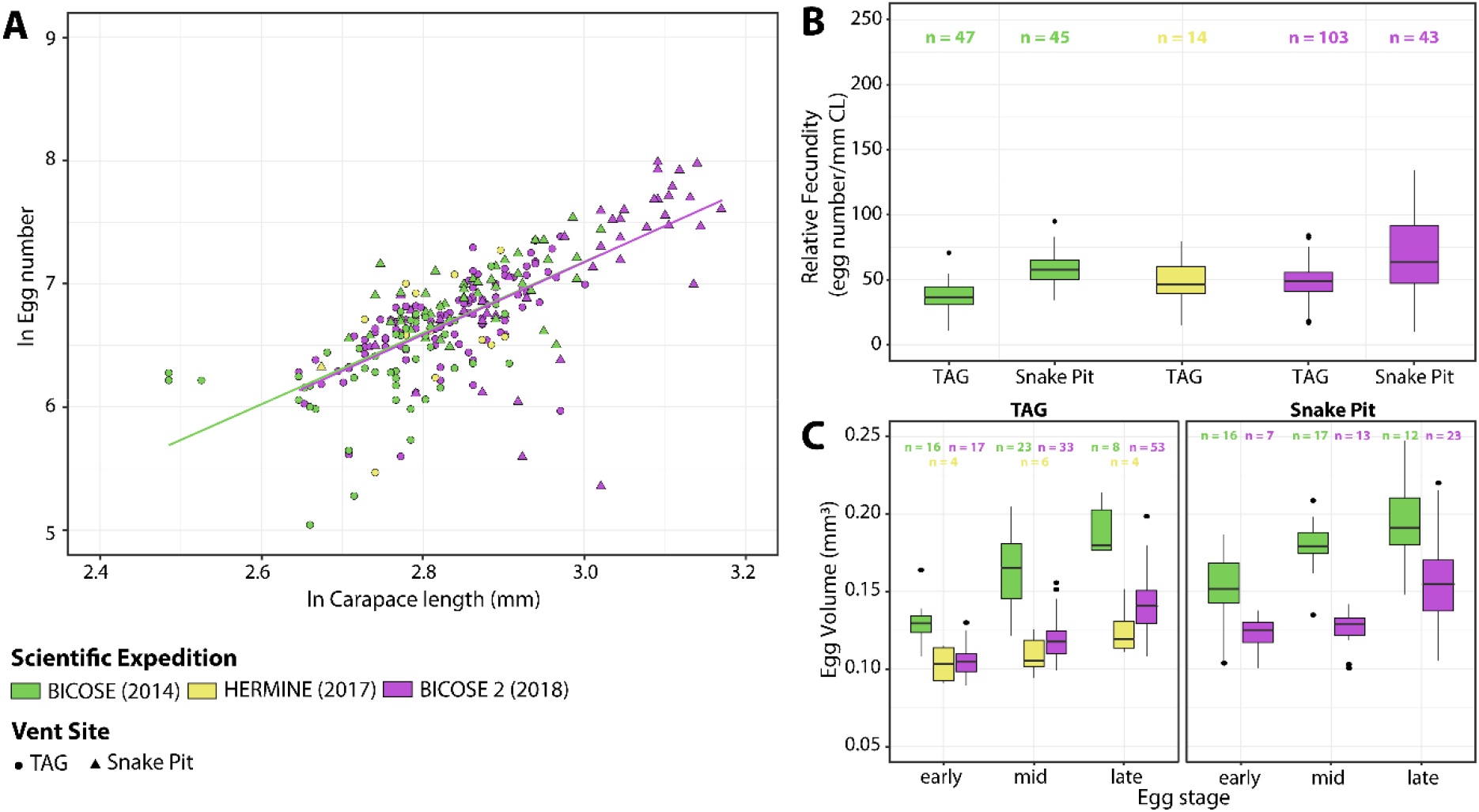
Interannual comparison of reproductive features of *Rimicaris exoculata* collected between 2014 and 2018. n: number of individuals for each condition, **a**. Variation of log_e_-transformed minimum realized fecundity with log_e_-transformed carapace length, **b**. Fecundity corrected for body size (carapace length) **c.** Egg volumes at different developmental stages.

Overall, size-specific fecundity ranged from 19 to 95 eggs mm^-1^ in 2014, from 15 to 80 eggs mm^-1^ in 2017 and from 10 to 134 eggs mm^-1^ in 2018 (figure 2b). Limited differences in size-specific fecundities were observed between different sampling years (Kruskal-Wallis, *H*= 65.025 *p* < 0.05) with significant variations between 2014 and 2018 at TAG only (2014: 37.3 ± 11.4; 2018: 49.4 ± 12.9; Dunn’s Multiple Comparison Test, *p* < 0.05). Differences in size-specific fecundity were also observed between the two vent fields both in 2014 and in 2018, (Dunn’s Multiple Comparison Tests, *p* < 0.05). Carapace length of brooding females from TAG were indeed larger in 2018 than in 2014 (Dunn’s Multiple Comparison Test, *p* < 0.05) suggesting that size-specific fecundity increase was attributable to this larger body size in 2018. Similarly, carapace lengths of brooding females were always larger at Snake Pit compared to TAG both in 2014 and in 2018 (Dunn’s Multiple Comparison Tests, *p* < 0.05).

Variations were also observed in egg volumes between different sampling years (Kruskal-Wallis, *H*= 87.434, *p* < 0.05) (figure 2c). Larger egg volumes were found in 2014 compared to other years of sampling for both vent fields and across all developmental stages (Dunn’s Multiple Comparison Tests, *p* < 0.05). We note that the 2014 dataset was produced separately in a previous study (See supplementary material) and may be biased by an observer effect.

### Variations in reproductive patterns between *Rimicaris* species

Like *R. exoculata,* the realized fecundity of *R. kairei* and *R. chacei* correlated positively with carapace length (Spearman’s correlation: *R. kairei: r*= 0.40, *p*= 0.0044; *R. chacei: r*= 0.70, *p*= 0.00066) (figure 3a). Moreover, the slopes of the relationship between log_*e*_-transformed realized fecundity and log_*e*_-transformed carapace length did not significantly differ between *R. exoculata* and its sister species *R. kairei (F*= 1.6168, *p*= 0.2054). However, they significantly differed between *R. exoculata* and its co-occurring congener *R. chacei* (*F*= 6.6376, *p* < 0.05).

**Figure 3.**
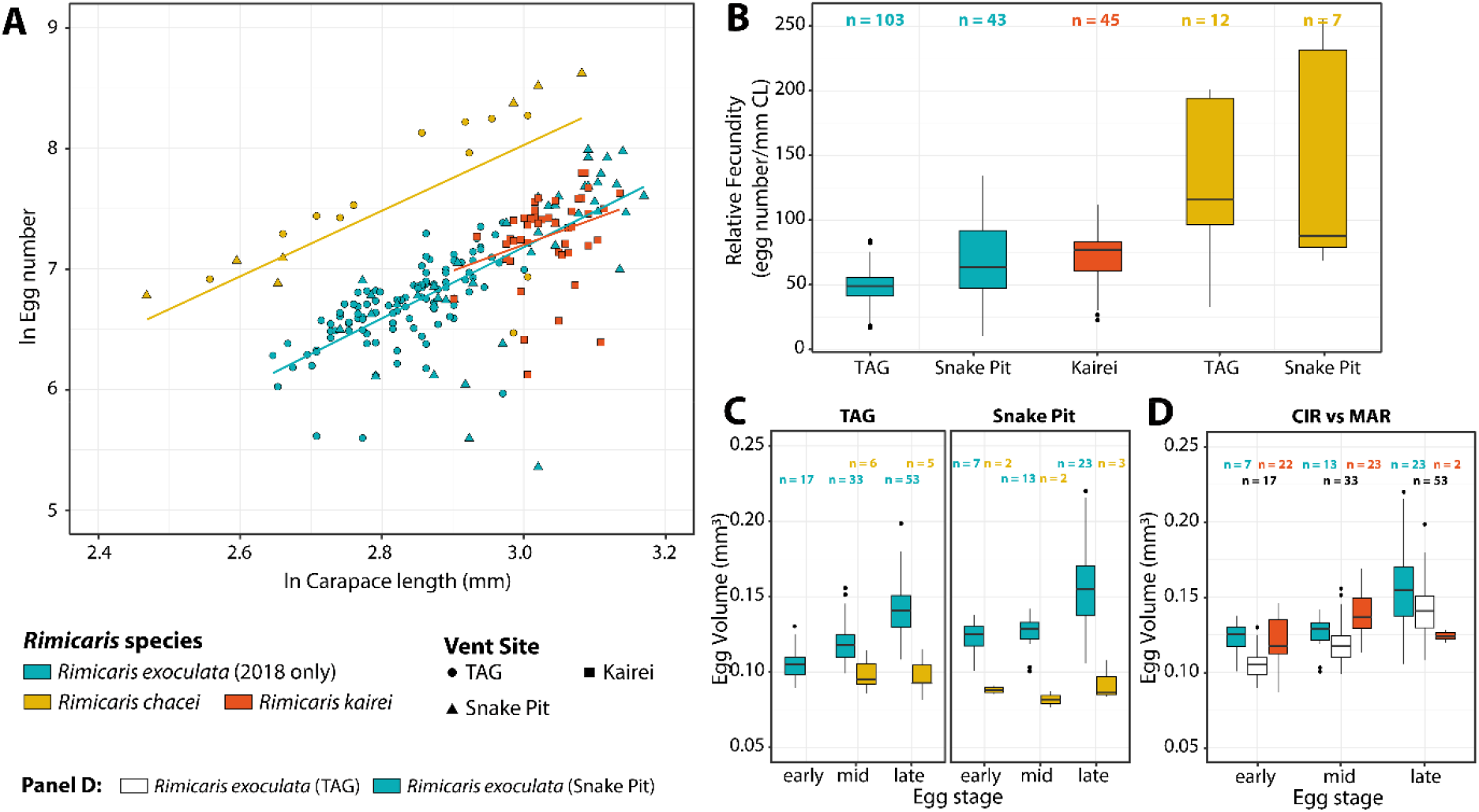
Comparison of reproductive features between *Rimicaris* species from the MAR and the CIR. n: number of individuals for each condition, **a.** Variation of log_e_-transformed minimum realized fecundity with log_e_-transformed carapace length, **b.** Fecundity corrected for body size (carapace length), **c.** Egg volumes at different developmental stages of *Rimicaris* shrimps from the MAR. **d.** Egg volumes at different developmental stages of *R. exoculata* and *R. kairei.*

Overall, size-specific fecundity ranged from 23 to 112 eggs mm^-1^ for *R. kairei* and from 26 to 255 eggs mm^-1^ for *R. chacei* (figure 3b). Large differences in size-specific fecundity were observed between *R. exoculata* and other *Rimicaris* species (Kruskal-Wallis *H*= 68.836, p < 0.05). Size-specific fecundity of *R. chacei* was always significantly greater compared to *R. exoculata* both at TAG *(R. exoculata:* 49 ± 13; *R. chacei:* 129 ± 59; Dunn’s Multiple Comparison Test, *p* < 0.05) and Snake Pit *(R. exoculata:* 69 ± 31; *R. chacei:* 148 ± 87; Dunn’s Multiple Comparison Test, *p* < 0.05). These differences were independent of size variations between the two species with *R. chacei* brooding females being either of similar size at TAG (Dunn’s Multiple Comparison Test, *p* > 0.05) or even smaller at Snake Pit (Dunn’s Multiple Comparison Test, *p* < 0.05). Less differences in size-specific fecundities were observed between *R. exoculata* and *R. kairei* with only a higher size-specific fecundity for *R. kairei* (72 ± 20) from Kairei compared to *R. exoculata* from TAG (49 ± 13; Dunn’s Multiple Comparison Test, *p* < 0.05). Such variations may be linked to differences in body size of brooding females, as *R. exoculata* from TAG were smaller in average than *R. kairei* from Kairei (Dunn’s Multiple Comparison Test, *p* < 0.05).

Large differences were also observed in egg volumes between the three *Rimicaris* species (Kruskal-Wallis *H=* 49.80, *p* < 0.05) (figure 3c,d). Thus, egg volumes of *R. chacei* were significantly smaller than those of *R. exoculata* at any stage across all vents (Dunn’s Multiple Comparison Test, *p* < 0.05) (figure 3c). In contrast, differences in egg volumes were more limited between *R. exoculata* and *R. kairei* with significantly larger eggs for *R. kairei* from Kairei compared to *R. exoculata* from TAG for mid stages only (figure 3d). Surprisingly, we did not find gradual increase of egg volumes between successive developmental stages of *R. chacei* as was observed in the two other *Rimicaris* species, either at TAG (Kruskal-Wallis *H*= 2.57, *p* > 0.05) or at Snake Pit (Kruskal-Wallis *H*= 0.61, *p* > 0.05). Of note, we did not observe ready-to-hatch or hatched egg broods in *R. chacei* as was seen in the other shrimps.

### Temporal variations in the presence of *Rimicaris* ovigerous females

A large number of ovigerous females of *R. exoculata* and *R. chacei* were retrieved in February 2018 at both MAR vent fields (figure 4a). Indeed, 55.4% and 18.2% of sexually mature *R. exoculata* females collected from TAG and Snake Pit respectively, were ovigerous females, while these account for 70.6% and 32.3% of sexually mature *R. chacei* females from TAG and Snake Pit, respectively. Ovigerous females of *R. exoculata* were also present between late March and April 2017 although in a relatively lower proportion, constituting 25.7% and 4% of sexually mature females collected at TAG and Snake Pit, respectively (figure 4a). The majority of *R. exoculata* ovigerous females were collected in dense aggregations close to vent orifices with only two ovigerous females taken outside these assemblages. On the other hand, ovigerous females of *R. chacei* were collected either in hidden aggregations where large populations of their adults have been retrieved (Methou et al. 2021) or within dense aggregations of *R. exoculata.* As highlighted previously (Hernández-Ávila et al. 2021), ovigerous females of *R. exoculata* were almost absent from samples collected outside this period of the year – i.e., January to April – despite more than 35 years of sampling efforts along the MAR (figure 4a). Similarly, ovigerous females of *R. chacei* were almost absent from other sampling reports with only two ovigerous females collected in January 2014 at Snake Pit and one in June 1994 at Lucky Strike (figure 4b)

**Figure 4.**
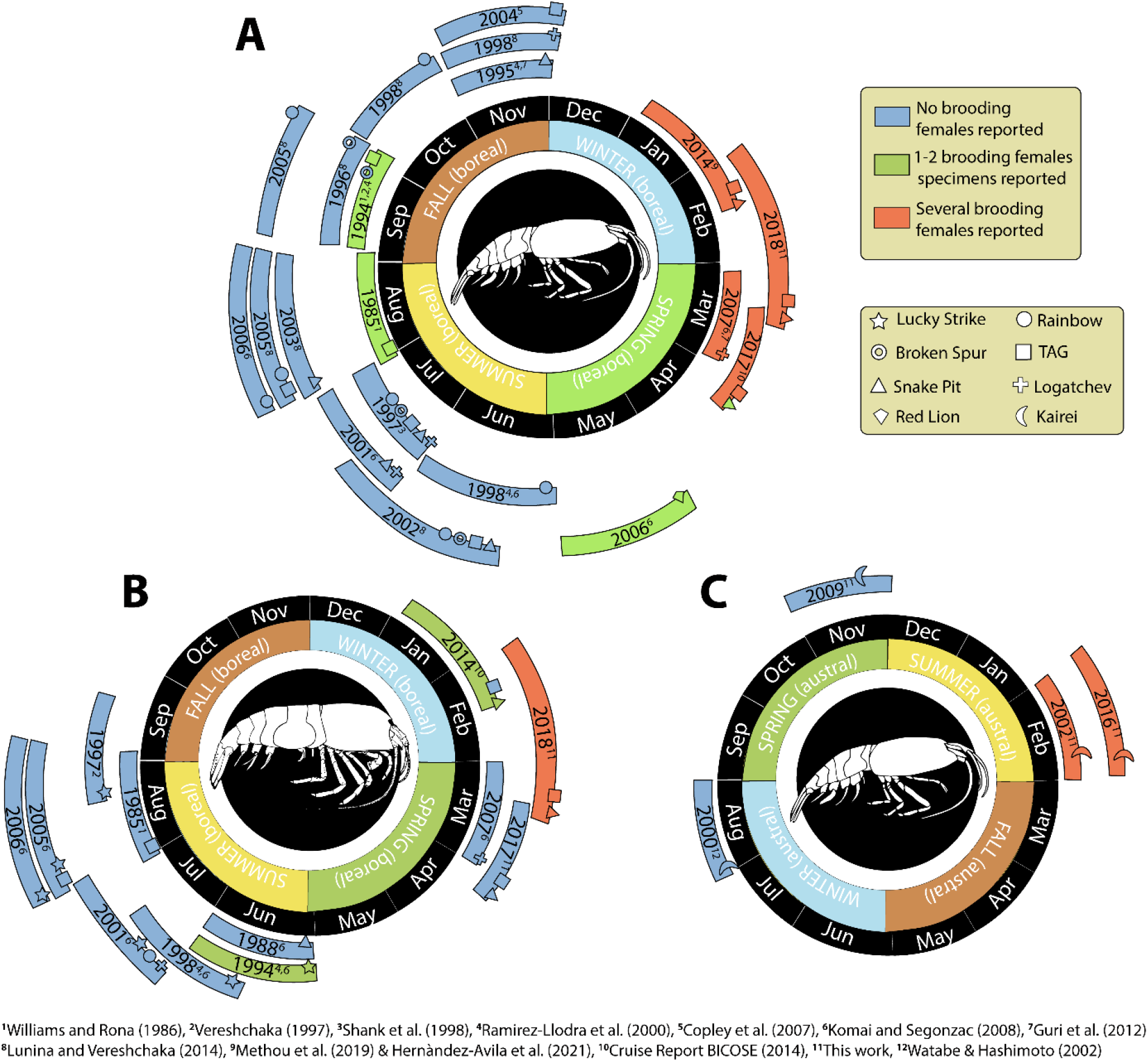
Diagrams summarizing occurrence of reproductive stages of the studied *Rimicaris* shrimp species over different samplings realized between 1985 and 2018 according to the year period **a.** for *R. exoculata.* **b.** for *R. chacei.* **c.** for *R. kairei.*

Large numbers of ovigerous *R. kairei* were also retrieved in February 2002 and 2016 at Kairei on the CIR (figure 4c), constituting respectively 16.3% and 19.2% of sexually mature females collected at this vent field. Conversely, ovigerous females were completely absent from samples collected in November 2009 (figure 4c). Like *R. exoculata,* all of these ovigerous females were collected in dense aggregations of shrimps next to hot fluid emissions.

## Discussion

### Distinct reproductive strategies in *Rimicarís* shrimps with evidence of diet influence

*Rimicaris exoculata* and *Rimicaris chacei* exhibit distinct reproductive traits unlikely to simply result from differences in mean body sizes. *Rimicaris kairei* is similar to its sister species *R. exoculata,* in terms of size-specific fecundity or mean egg size. Our results indicate that the two lineages (*R. exoculata* and *R. kairei,* vs *R. chacei)* are situated at two ends of the trade-off between size-specific fecundity and egg sizes currently observed within alvinocaridids (Ramirez-Llodra et al. 2000, Ramirez-Llodra and Segonzac 2006, Copley and Young 2006, Nye et al. 2013). On one hand, *R. chacei* brooding females display one of the highest size-specific fecundity and one of the smallest mean egg volumes among alvinocaridids, whereas *R. exoculata/R. kairei* brooding females present a rather low size-specific fecundity and a large mean egg volume for this family.

These variations mirror differences observed in benthic post-settlement stages of these two lineages, with significantly smaller body size for settled juveniles, as well as much larger relative proportions of juveniles in populations of *R. chacei* (Methou et al. 2020, 2021). Higher egg numbers per brood in *R. chacei* might thus contribute to its high supply of settlers despite an apparently lower breeding population size than *R. exoculata* in northern MAR vent fields. Previous work has hypothesized distinct larval life histories between these two species based on distinct stable isotope ratios (Methou et al. 2020). Given we did not observe ready-to-hatch or hatched egg broods for *R. chacei,* it is unclear if observed differences in mean egg size between the two species would ultimately lead to distinct larval sizes at hatching. No size variation was found among first zoeal larval stages of four phylogenetically distinct alvinocaridids (Hernández-Ávila et al. 2015). Although *R. chacei* zoea have not been observed so far, it is unlikely that hatching size difference only, if it exists, explains the large size difference at recruitment between *R. exoculata* and *R. chacei,* which also probably results from differences in their pelagic lives during their dispersal phase.

The sister species of *R. chacei* is *R. hybisae* from vents on the Mid Cayman Spreading Centre, but interestingly size-specific fecundity of *R. hybisae* is more similar to *R. exoculata/R. kairei* rather than *R. chacei* (Nye et al. 2013). *Rimicaris hybisae* share with *R. exoculata/R. kairei* both a comparable ecological context, living in dense aggregations close to vent fluids emissions, and a comparable gross morphology, with an enlarged cephalothorax (Zbinden and Cambon-Bonavita 2020), characters lacking in *R. chacei.* These are linked to distinct feeding strategies between the two groups of shrimps, with a mixotrophic diet for *R. chacei* (Gebruk et al. 2000, Apremont et al. 2018, Methou et al. 2020) and dependency on chemosynthetic symbionts for others including *R. hybisae* (Rieley et al. 1999, Gebruk et al. 2000, Van Dover 2002, Streit et al. 2015, Methou et al. 2020). Variations in fecundity according to feeding strategies have been widely recognized in marine animals such as between carnivorous and herbivorous sea turtles (Broderick et al. 2003), among seabirds with different feeding habits (Pierotti and Annett 1990) or between *Capitella* worms fed with different food sources in laboratory (Qian and Chia 1991). Reduced or increased fecundities have also been observed experimentally in several shrimp species according to their diets (Lin and Shi 2002, Tziouveli et al. 2011, Almeida et al. 2018). We therefore posit that the differences we observe between fecundities in our three *Rimicaris* shrimps should be linked to their feeding ecology, rather than being constrained by their phylogeny.

Despite these variations in average egg number, mean egg sizes of *R. hybisae* and *R. chacei* were in contrary comparable with similar maximum and minimum egg diameters (Nye et al. 2013). As *R. exoculata* and *R. kairei* also exhibit similar egg volumes, phylogenetic constraints are possibly exerted on mean egg sizes for these shrimps. Further work on more alvinocaridid shrimps with different feeding strategies should help to elucidate these questions.

### The intriguing reproductive cycles of *Rimicaris* shrimps

We report here a much-increased presence of ovigerous *R. exoculata* in February 2018 and to a lower extent in late March-April 2017, compared to other year periods. Combined with previous data from the literature (summarized in figure 4A), these results support the existence of a brooding period for *R. exoculata* mostly during boreal winter, starting in January/February and ending around March/April. Moreover, we did not observe any variability in reproductive features of *R. exoculata,* either in terms of fecundity or proportions of reproductive females, between different sampling years – indicating a relative inter-annual stability of these traits (figure 2). Since gamete production can be sustained continuously in species depending on local chemosynthesis for their diet, mechanisms driving periodicity in shrimp life cycles are more likely to act on their planktotrophic larval phase through seasonality in availability of photosynthetically derived food. The few previous studies looking at a limited number of samples indeed reported asynchronous ovarian development for *R. exoculata* suggesting continuous egg production (Ramirez-Llodra et al. 2000, Copley et al. 2007). However, the oocyte maximum size appeared to be lower in individuals collected summer than in those collected in autumn, regardless of the vent site origin [16, 34], which may also point at a gradual accumulation of mature oocytes in ovaries, culminating in winter, prior to an increase in females spawning activity. Additional observations of intraovarian development through the year are clearly needed to better characterize rhythms in oocyte production, which may also well be decorrelated from spawning activity as seen in some deep-sea corals from the MAR (Rakka et al. 2021). Large proportions of *R. chacei* ovigerous females were also retrieved in February 2018 at the two vent fields, in contrast to historical sampling at other periods along the MAR (figure 4B). Hence, a similar periodicity in reproduction likely exists for both *Rimicaris* shrimps at MAR. Ovigerous *R. chacei* females were however scarce in other expeditions both during and outside this period (figure 4B), which could be due to difficulties in sampling this rather fast swimming species, thus preventing so far, a full appreciation of its reproductive activity.

We also found large proportions of *Rimicaris kairei* ovigerous females during austral summer in February 2002 and 2016, while they were absent from collections in August 2000 and November 2009 (figure 4c). Spawning in *R. kairei* thus appears to follow the same timing as *R. exoculata* and *R. chacei.* Such pattern was unexpected given their distribution in different hemispheres, at vent fields with opposite latitudes (figure 1). Reproductive periodicity in deepsea animals has often been related to seasonal variations in surface photosynthetic production [26]-[28], and *Rimicaris* shrimps from the two hemispheres are expected to experience opposite trends in terms of surface primary production along the year (Pabortsava et al. 2017, Harms et al. 2021). driving opposite reoroductive timing. Although our dataset for *R. kairei* is more limited than for *R. exoculata,* these sister species thus appear to share same periodic brooding cycle, regardless of local seasonal variations in surface photosynthetic productivity. *Rimicaris* species studied here provide a first example of reproductive periodicity in deep-sea ecosystems apparently unlinked to seasonal variations of photosynthetic materials from the surface.

Besides light and food supply, temperature has also been recognized as an important external time-giving cue in marine organisms (Mat 2019). Compared to most of the deep sea where temperature is relatively constant within regions, hydrothermal vents constitute heat anomalies with steep and unpredictable gradients at small spatial scales, even within a same animal aggregation (Schmidt et al. 2008). Thermal variability at an annual scale could perhaps serve as a time-synchronising cue for these *Rimicaris* shrimps. So far, thermal variations in these ecosystems at a year-round scale remain poorly documented. However, long-term observatories deployed on two vent fields from the northern hemisphere have reported clear tidal influence, although temperatures are relatively stable throughout the year (Barreyre et al. 2014, Cuvelier et al. 2017). Recently, it was revealed that bathymodiolin mussels inhabiting vents on the MAR retain and express circadian clock genes following tidal cycles which may be mediated by either tidal stimulus or internal clock (Mat et al. 2020). Possibly, *Rimicaris* shrimp species studied herein also use a set of biological clocks to time their reproductive activity. Though, not all alvinocaridid shrimps seem to exhibit periodicity in their reproductive patterns, despite a gametogenesis phase being strongly constrained phylogenetically (Ramirez-Llodra et al. 2000). For example, regular sampling of *Mirocaris fortunata* yielded ovigerous females throughout the year (Ramirez-Llodra et al. 2000, Komai and Segonzac 2003, Methou 2019) and similarly in *Rimicaris variabilis* (Komai and Tsuchida 2015, Komai et al. 2016). It therefore appears unlikely that phylogenetic constraints alone have maintained a periodic brooding in several alvinocaridid genera but not in others. Then again, other alvinocaridid species may also indeed follow seasonality in surface production, like *Alvinocaris stactophila* with a brooding period between November and March (Copley and Young 2006), and inhabiting relatively shallow cold seeps under highly productive coastal waters. Hence, several synchronizing factors may co-exist in vents and other deep-sea ecosystems, even within the same family of animals.

For example, our results here demonstrate that *R. exoculata* and *R. kairei* exhibit seasonality decoupled from surface seasonality, but as the study areas have relatively low surface primary productivity and export of particulate organic carbon (POC) to the deep (Pabortsava et al. 2017, Harms et al. 2021) it is also possible that a higher POC export may begin to interfere with this pattern. Taken together – primary surface production may be the key synchronising cue for some deep-sea species as demonstrated in bathymodiolin mussels and vent crabs (Dixon et al. 2006, Tyler et al. 2007), but our results underscore that reproductive cycles and seasonality at the deep is not as simple as it was thought to be.

A thorough understanding of reproductive ecology of the dominant species shaping the ecosystem is key to assessing the resilience of deep-sea ecosystems, but little has been done so far even for emblematic taxa like *Rimicaris.* With deep-sea mining for massive sulphides and other resources being imminent (Van Dover et al. 2018), the habitat of these amazing animals that can live nowhere else is being threatened. Elucidating the ecology of these inhabitants of the deep is now more important than ever, and our data exemplifies the importance of time-series studies. Research expeditions in a particular region tend to be conducted in similar timings of the year due to seasonal variations in sea and weather conditions, but our results show that communities need to be revisited at different times of the year.

### Ethics

The animals used in this study were invertebrate carideans collected in the French contract area for the exploration of polymetallic sulphur deposits (Mid-Atlantic Ridge) or International Waters (Central Indian Ridge). Each individual was immediately preserved in ethanol after recovery and no live experiments were carried out.

## Author’s contributions

P.M. collected and sorted specimens in 2018 on the MAR, carried out the identification and measurement of specimens from all sampling cruises, lead the data analysis, participated in the conception and design of the study and drafted the manuscript; C.C. carried out the collection of specimens in 2016 on the CIR and helped draft the manuscript; H.K.W. carried out the collection of specimens in 2009 and critically revised the manuscript; M-A.C-B. collected specimens in 2017 on the MAR, coordinated the study and critically revised the manuscript; F.P. collected and sorted specimens in 2017 conceived and designed the study, coordinated the study and helped draft the manuscript. All authors gave final approval for publication and agreed to be held accountable for the work performed therein. The authors declare they have no competing interests.

## Acknowledgements

The authors thank the captains and crews of R/V *Pourquoi pas?* and HOV *Nautile* submersible team for their efficiency, as well as the chief scientists and scientific parties of HERMINE (chief scientist: Yves Fouquet; http://dx.doi.org/10.17600/17000200) and BICOSE 2 cruise (chief scientist: Marie-Anne Cambon-Bonavita; http://dx.doi.org/10.17600/18000004). Further thanks also go to the captains and crews of R/V *Yokosuka,* and the HOV *Shinkai 6500* submersible team for their continuous support of the scientific activity at sea, as well as the chief scientists and scientific parties during the research cruises YK01-15 (chief scientist: Ken Takai), YK09-13 (chief scientists: Kentaro Nakamura, Satoshi Nakagawa) and YK16-E02 (chief scientist: Ken Takai). We also thanks Dr. Ivan Hernández-Ávila (Facultad de Ciencias Naturales, Universidad Autónoma del Carmen) for his help to sort the specimens on board and the analyses to determine sexual maturity of these shrimps.

