## supplemental_material.pdf for "Reproduction in deep-sea vent shrimps is shaped by diet and phylogeny, with rhythms unlinked to surface production"

### Materials & Methods

#### Estimation of the size of effective sexual maturity (ESM)

The size of effective sexual maturity (ESM) of *R. chacei* and *R. kairei* was estimated according to (Hernández-Ávila et al., 2021; King, 2007), from the proportion of ovigerous females per size class, corrected by the maximum proportion of ovigerous females.

The proportion of ovigerous female specimens were plotted against size classes and fitted to the logistic equation: $Povf = 1/(1+e^{(a-b*CL)})$ where Povf is the proportion of ovigerous shrimp females, CL is the carapace length, and a and b are constants. The ESM was estimated by the size at which Povf is 50% (figure S1).

### Results

#### Observer effect

A sub-sample of the broods from 2014 were measured again by the same observer who produced the 2017 and 2018 datasets, and compared with the dataset produced by the first observer. Accordingly, large differences were found between the first dataset from 2014 and the re-examined sub-sample but not between samples treated by the same observer (figure S2).

#### Developmental stages proportions

Proportions of each developmental stage were rather similar between 2014 and 2017 (TAG: $\chi^2 = 0.546$, 2 df, $p > 0.05$) with a majority of mid-stage egg broods (2014: 48.9 %; 2017: 42.9 %) (figure S3a). In contrast, these proportions differ between 2014 and 2018 at both vent fields (TAG: $\chi^2 = 27.938$, 2 df, $p < 0.05$; Snake Pit: $\chi^2 = 58.493$, 2 df, $p < 0.05$) as most egg broods from 2018 were at a late developmental stage (TAG: 51.5%; Snake Pit: 53.5%) (figure S3a).

In addition, proportions of each developmental stages were relatively similar between *R. exoculata* and *R. chacei* broods at both sites (TAG: $\chi^2 = 84.837$, 2 df, $p < 0.05$; Snake Pit: $\chi^2 = 84.837$, 2 df, $p < 0.05$) with a majority of late-stage egg broods (figure S3b). On the other hand, *R. kairei* egg broods from Kairei were for most of them at an early or mid-developmental stage (early: 46.8%; mid: 48.9%).

#### Estimation of the size of effective Sexual Maturity (ESM)

We estimate the size at effective sexual maturity (ESM) at 12.65 mm for *R. chacei* females (figure 1a) and at 15.4 mm for *R. kairei* (figure 1b). Thus, females with size ≥ 12.65 mm for *R. chacei* and with size ≥ 15.4 mm for *R. kairei* were considered as sexually mature.

These ESM sizes correspond respectively to 58% and 65% of the maximal size reported for each species (*R. chacei*: 21.8 mm; *R. kairei*: 23.7 mm). Additionally, we reevaluate this proportion given previously for *R. exoculata* (Hernández-Ávila et al., 2021), which ESM represent 62% of the new maximal size reported for this shrimp species in our 2018 dataset (*R. exoculata*: 24.4 mm ;(Methou et al., 2021)).

#### Spatial variations in the presence of *Rimicaris* ovigerous females

Important spatial variations in the proportions of ovigerous females could be observed between the different dense aggregate assemblages. These were comprised between 5.6% and 69.6%

for those collected in 2018 (figure S4) and between 3.6% and 21.6% of the sexually mature females from assemblages collected in 2017 (figure S5). Additionally, ovigerous females were absent in 4 out of the 7 dense aggregates collected in 2017 but only in 3 out of the 15 dense aggregates collected in 2018.

Spatial variations in the proportions of ovigerous females among samples were also observed for *R. kairei* and *R. chacei*. These were comprised between 3.4% and 24.1% of sexually mature *R. kairei* females in February 2016 (figure S6) and between 46.2% and 73.3% of sexually mature *R. chacei* females in February 2018 (figure S4). These reproductive stages were present in every dense aggregate of *R. kairei* and three out of the four populations of *R. chacei* where the number of adults exceed five individuals.

#### Figure Caption

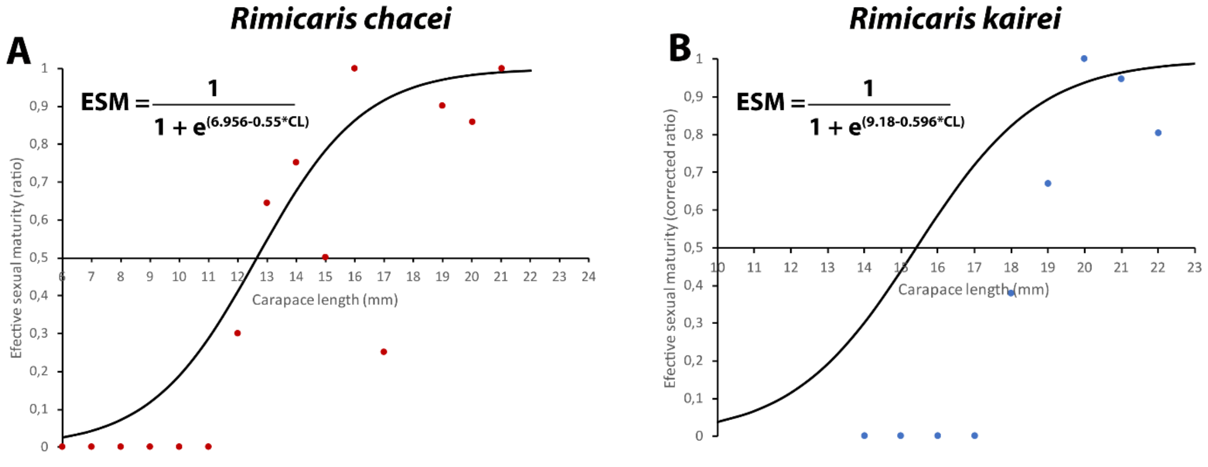


**Figure S1.** Estimation of effective sexual maturity (ESM) of **a.** *R. chacei* and **b.** *R. kairei*.

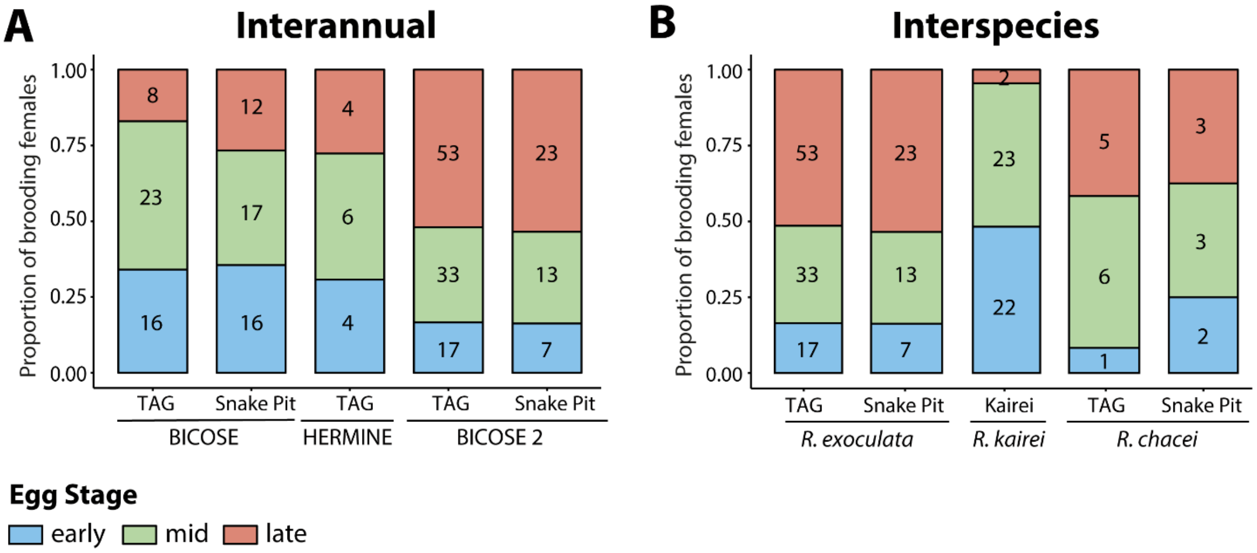


**Figure S2.** Proportions of each egg developmental stages. Numbers within the plots indicates the number of broods per developmental stages. **a.** Comparison of egg developmental stage proportions between *R. exoculata* broods from different sampling years. **b.** Comparison of egg developmental stage proportions between broods of the different *Rimicaris* species from the MAR and the CIR.

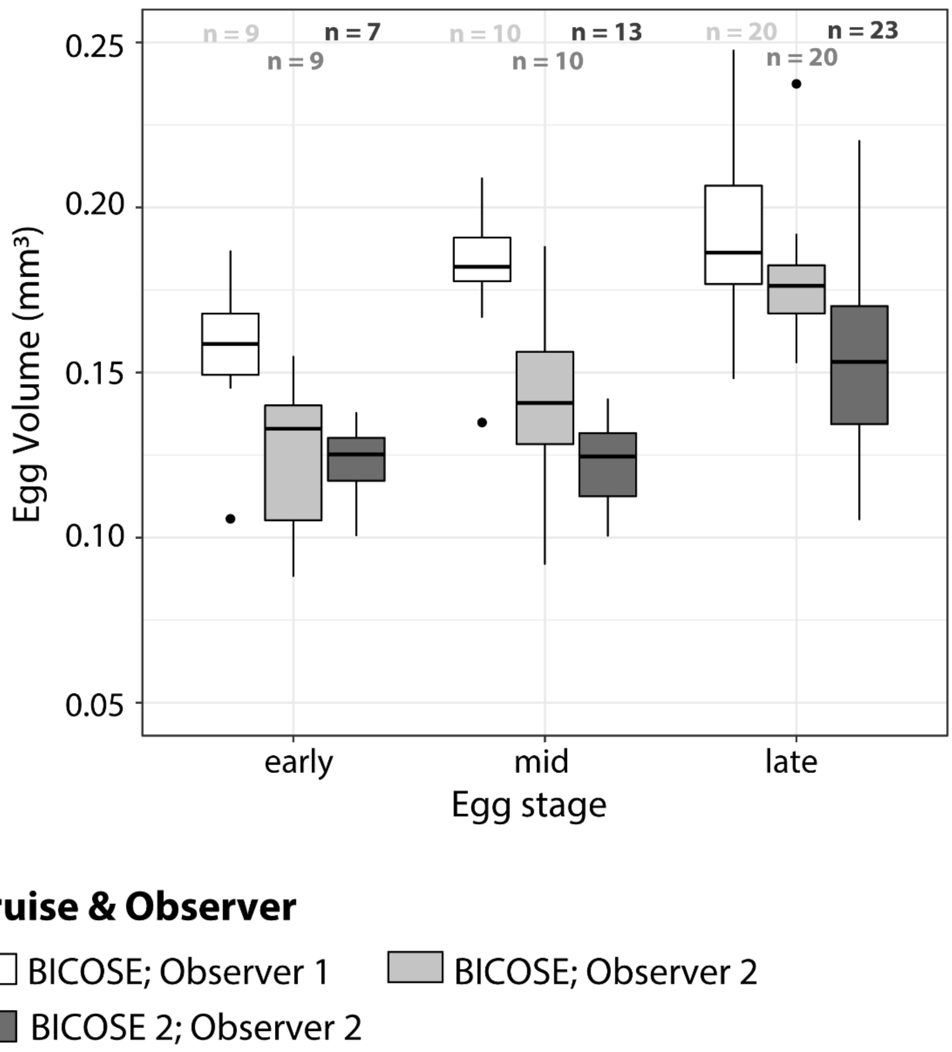


**Figure S3.** Evaluation of a potential observer effect in the comparison of egg volumes collected from different sampling expeditions.

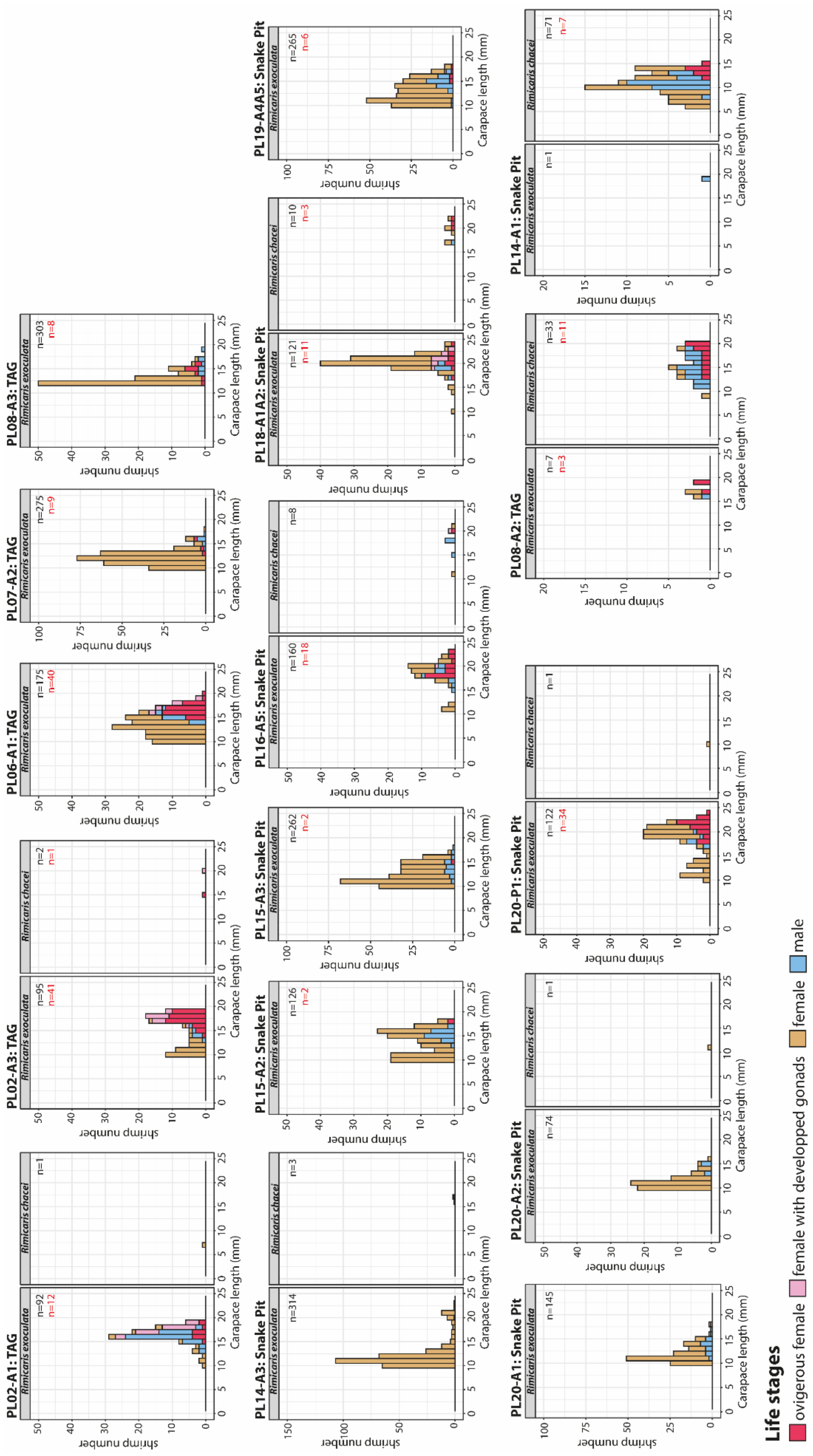


**Figure S4.** Small-scale spatial variations in size-frequency distribution of *Rimicaris* reproductive stages collected during the BICOSE 2 expedition in 2018. **n:** (black) number of measured adult individuals; (red) number of measured ovigerous females.

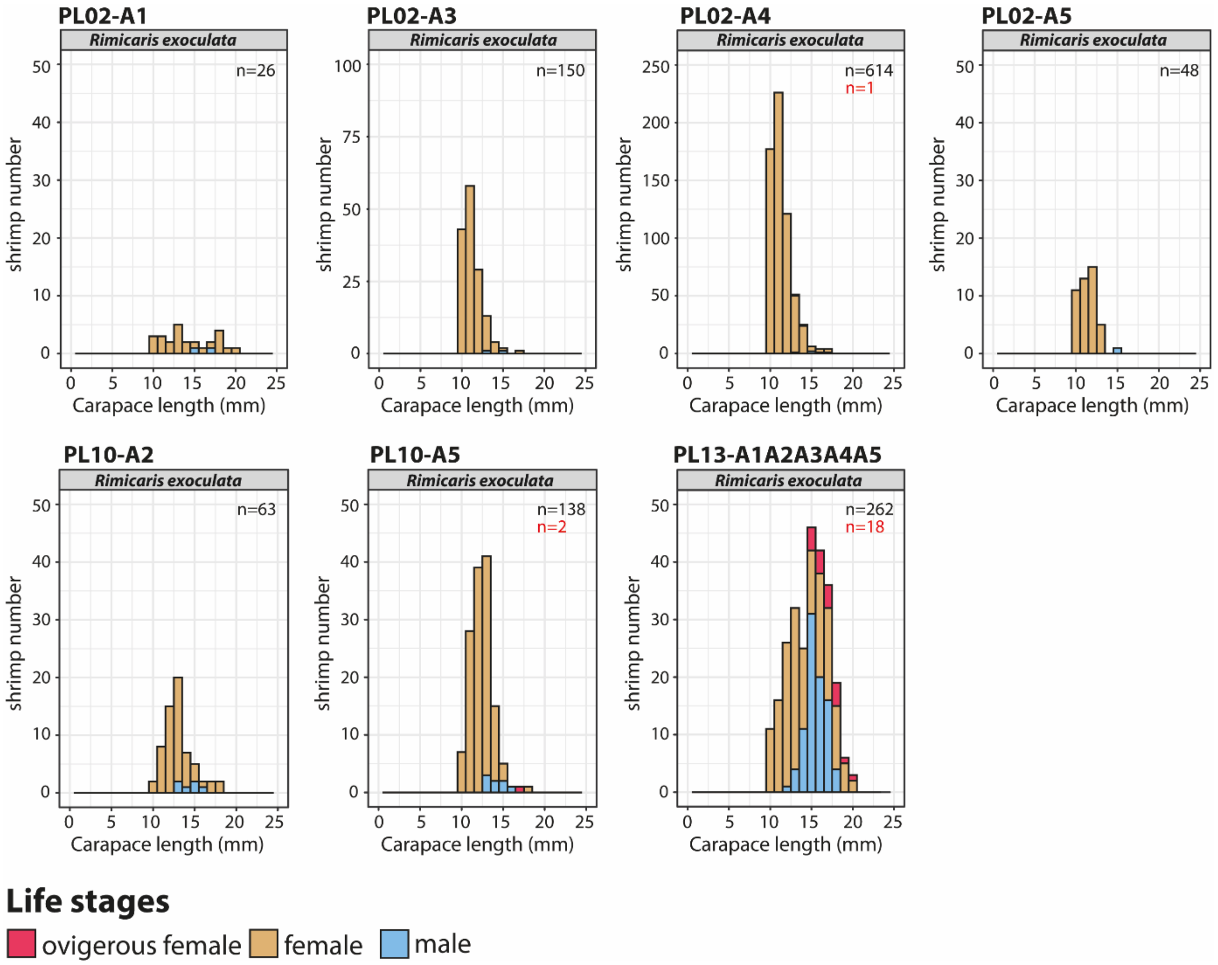


**Figure S5.** Small-scale spatial variations in size-frequency distribution of *R. exoculata* reproductive stages collected during the HERMINE expedition in 2017. **n:** (black) number of measured adult individuals; (red) number of measured ovigerous females.

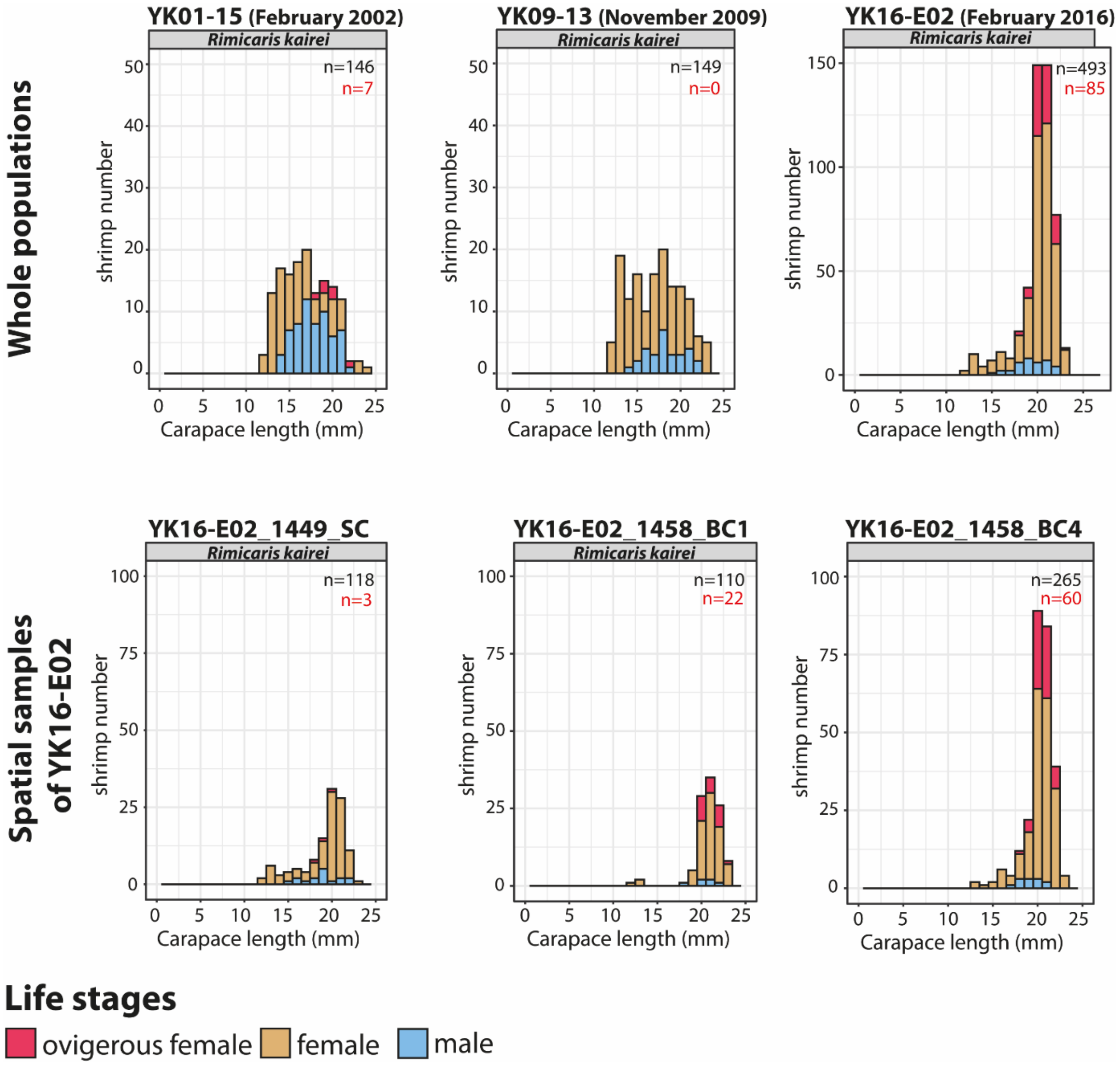


**Figure S6.** Small-scale spatial variations in size-frequency distribution of *R. kairei* reproductive stages collected during the YK16-E02 2 expedition in 2016. **n:** (black) number of measured adult individuals; (red) number of measured ovigerous females.

| **A. Relative Fecundity** | BICOSE TAG | BICOSE Snake Pit | HERMINE TAG | BICOSE 2 TAG |
|---|---|---|---|---|
| BICOSE TAG | | | | |
| BICOSE Snake Pit | **1.47e-10** | | | |
| HERMINE TAG | 3.69e-02 | 5.97e-02 | | |
| BICOSE 2 TAG | **3.14e-05** | **2.16e-03** | 9.14e-01 | |
| BICOSE 2 Snake Pit | **6.51e-11** | 8.89e-01 | 4.17e-02 | **9.07e-04** |

| **B. Carapace length (mm)** | BICOSE TAG | BICOSE Snake Pit | HERMINE TAG | BICOSE 2 TAG |
|---|---|---|---|---|
| BICOSE TAG | | | | |
| BICOSE Snake Pit | **6.43e-06** | | | |
| HERMINE TAG | 1.07e-01 | 1.99e-01 | | |
| BICOSE 2 TAG | **1.08e-04** | 1.88e-01 | 5.83e-01 | |
| BICOSE 2 Snake Pit | **5.32e-17** | **2.76e-04** | **1.36e-04** | **1.09e-08** |

| **Egg volume ($mm^3$) Early stage** | BICOSE TAG | BICOSE Snake Pit | HERMINE TAG | BICOSE 2 TAG |
|---|---|---|---|---|
| BICOSE TAG | | | | |
| BICOSE Snake Pit | 9.45e-02 | | | |
| HERMINE TAG | **4.06e-02** | **1.81e-03** | | |
| BICOSE 2 TAG | **1.77e-03** | **9.88e-07** | 8.05e-01 | |
| BICOSE 2 Snake Pit | 4.92e-01 | 5.55e-02 | 1.40e-01 | 9.31e-02 |

| **Egg volume ($mm^3$) Mid stage** | BICOSE TAG | BICOSE Snake Pit | HERMINE TAG | BICOSE 2 TAG |
|---|---|---|---|---|
| BICOSE TAG | | | | |
| BICOSE Snake Pit | 2.41e-01 | | | |
| HERMINE TAG | **2.08e-04** | **6.60e-06** | | |
| BICOSE 2 TAG | **1.61e-06** | **7.41e-09** | 4.02e-01 | |
| BICOSE 2 Snake Pit | **9.89e-03** | **2.54e-04** | 1.61e-01 | 2.89e-01 |

| **Egg volume ($mm^3$) Late stage** | BICOSE TAG | BICOSE Snake Pit | HERMINE TAG | BICOSE 2 TAG |
|---|---|---|---|---|
| BICOSE TAG | | | | |
| BICOSE Snake Pit | 9.77e-01 | | | |
| HERMINE TAG | **4.93e-04** | **2.39e-04** | | |
| BICOSE 2 TAG | **8.25e-05** | **2.59e-06** | 2.37e-01 | |
| BICOSE 2 Snake Pit | **1.95e-02** | **6.52e-03** | **2.70e-02** | **2.34e-02** |

**Table S1.** Specific p-values of Dunn tests Interannual multiple comparisons of *Rimicaris exoculata*

| **A. Relative Fecundity** | *R. exoculata* TAG | *R. exoculata* Snake Pit | *R. chacei* TAG | *R. chacei* Snake Pit |
|---|---|---|---|---|
| *R. exoculata* TAG | | | | |
| *R. exoculata* Snake Pit | **1.30e-04** | | | |
| *R. chacei* TAG | **7.83e-07** | **2.30e-02** | | |
| *R. chacei* Snake Pit | **4.29e-05** | **3.88e-02** | 9.32e-01 | |
| *R. kairei* Kairei | **8.43e-08** | 2.91e-01 | 1.39e-01 | 1.31e-01 |

| **B. Carapace length (mm)** | *R. exoculata* TAG | *R. exoculata* Snake Pit | *R. chacei* TAG | *R. chacei* Snake Pit |
|---|---|---|---|---|
| *R. exoculata* TAG | | | | |
| *R. exoculata* Snake Pit | **1.52e-10** | | | |
| *R. chacei* TAG | 7.53e-01 | **4.52e-03** | | |
| *R. chacei* Snake Pit | 9.11e-01 | **1.26e-02** | 9.48e-01 | |
| *R. kairei* Kairei | **8.77e-19** | 1.34e-01 | **4.98e-05** | **5.60e-04** |

| **Egg volume (mm³) Early stage** | *R. exoculata* TAG | *R. exoculata* Snake Pit | *R. chacei* TAG | *R. chacei* Snake Pit |
|---|---|---|---|---|
| *R. exoculata* TAG | | | | |
| *R. exoculata* Snake Pit | 0.037 | | | |
| *R. chacei* TAG | - | - | | |
| *R. chacei* Snake Pit | - | - | - | |
| *R. kairei* Kairei | 0.054 | 0.724 | - | - |

| **Egg volume (mm³) Mid stage** | *R. exoculata* TAG | *R. exoculata* Snake Pit | *R. chacei* TAG | *R. chacei* Snake Pit |
|---|---|---|---|---|
| *R. exoculata* TAG | | | | |
| *R. exoculata* Snake Pit | 1.76e-01 | | | |
| *R. chacei* TAG | **4.58e-02** | **6.57e-03** | | |
| *R. chacei* Snake Pit | - | - | - | |
| *R. kairei* Kairei | **4.15e-04** | 1.59e-01 | **4.45e-05** | - |

| **Egg volume (mm³) Late stage** | *R. exoculata* TAG | *R. exoculata* Snake Pit | *R. chacei* TAG | *R. chacei* Snake Pit |
|---|---|---|---|---|
| *R. exoculata* TAG | | | | |
| *R. exoculata* Snake Pit | **0.039** | | | |
| *R. chacei* TAG | **0.004** | **0.019** | | |
| *R. chacei* Snake Pit | **0.0002** | **0.002** | 0.936 | |
| *R. kairei* Kairei | - | - | - | - |

**Table S2.** Specific p-values of Dunn tests multiple comparisons between *Rimicaris* species
